# The *Drosophila* ovarian terminal filament imports lipophilic molecules that support cyst and follicle development within the ovariole

**DOI:** 10.1101/2025.07.30.667757

**Authors:** Bhawana Maurya, Allan C Spradling

## Abstract

Oogenesis in *Drosophila* requires small molecules as biosynthetic precursors and regulators of follicle development. But how such molecules reach germline stem cells (GSCs), developing germline cysts and follicles in the germarium and early ovariole is poorly understood. The flat, stacked cells of the terminal filament (TF) form a specialized somatic structure positioned at the anterior end of ovarioles in virtually all insect ovaries, but physiological roles TFs play in adult ovaries remain little known. By briefly knocking down, specifically in the TF, exocyst components affecting vesicle trafficking, lipid importers such as LpR2, and organic anion importers such as Oatp30B, we found that the TF provides lipophilic molecules to GSCs and downstream germarium cells needed to maintain lipid droplets, and germ cell differentiation.

When exocyst component Sec6 is knocked down, vesicles containing lipophilic cargos back up at the TF-germ cell junction, suggesting that endosomes move between the stacked TF cells and into cap cells by transcytosis. Some of these may be shuttled during the GSC cell cycle between the TF and newly forming fusome. Our studies suggest that TFs import lipophilic precursors and regulators to autonomously coordinate their ovariole’s stem cell activity and cyst development.

## INTRODUCTION

Ovaries in *Drosophila*, like most insect species, comprise a semi-autonomous collection of ovarioles, each containing germline cysts and ovarian follicles lined up in developmental order (reviewed in King, 1970; Büning, 1994). Characteristically, each ovariole starts with a "terminal filament (TF)," described in the earliest study of an insect ovary by Malphigi in 1669, but whose functional significance in adult egg production still remains unclear (Figure 1A). TF cells are flattened like a stack of coins, and their number varies between species (Sarikaya et al. 2011) and changes during adulthood (Forbes et al. 1996). The well studied *Drosophila* TF contains 8-9 somatic cells at adult eclosion that protrude from the end of each ovariole anteriorly, while posterior TF cells contact the GSC niche’s cap cells (Figure 1A).

**Figure 1:**
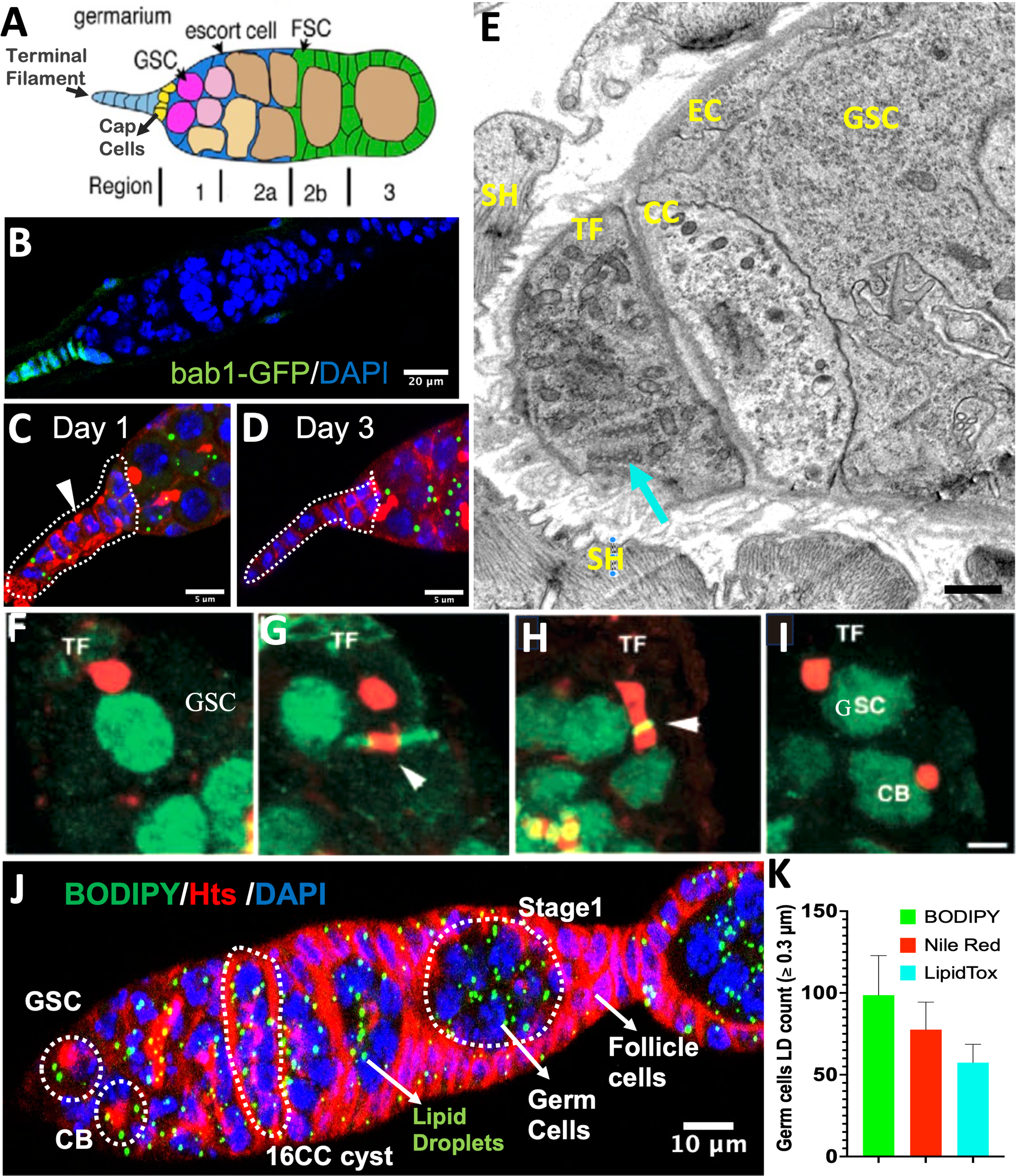
Identification of germ cell lipid droplets and Hts-positive aggregates in the terminal filament of the germarium. **(A)** Schematic diagram of the germarium, highlighting the terminal filament (TF), cap cells (cap) and anterior escort (EC) cells that maintain a niche for 2-3 germline stem cells (GSCs). Two follicle stem cells (FSCs) reside singly in separate small niches maintained by posterior ECs on each side of the germarium at the junction or region 2a and 2b, two of four germarium regions. **(B)** Germarium tip from a *bab1-GAL4* driven *nls:GFP* ovary stained with GFP and DAPI which marks the TF and CC cells. **(C)** Hts immunostaining reveals Hts-positive aggregates (triangle) within TF cells on day 1 post-eclosion. **(D)** Decline and disappearance of Hts aggregates in the TF by day3 of adulthood **(E)** An EM showing a rare cross section through a cap cell and the thin cytoplasm of a posterior TF cell, reveals the presence of abundant small vesicles (arrow). bar: 2μm. SH sheath, EC escort cell, GSC germline stem cell. **(F)** Image showing anterior anchoring of the GSC spectrosome at the TF/CC interface. **F-I** are from de Cuevas and Spradling, 1998. **(G)** Fusome plug formation at the ring canal in the cystoblast (CB). **(H)** Representative image showing fusion of the GSC spectrosome with the fusome plug in the cystoblast. **(I)** Round fusome establishment in the CB following fusion of the GSC spectrosome with the fusome plug, marking the completion of asymmetric division and specification of CB identity. **(J)** Lipid Droplets (LDs) were observed throughout the germarium using BODIPY493/503 staining. GSC, CB, 16CC **(K)** Quantification of lipid droplet number in germ cells of the germarium, stained with BODIPY, Nile Red (see Fig S1B) and LipidTox (see Fig S1C). Data are shown as mean ± SD; n = 52 germarium examined.

Terminal filaments form in early third instar (L3) larval ovaries when precursor cells intercalate in the nascent ovariole (Godt and Laski, 1995; Sahut-Barnola et al. 1995). TF precursors express two similar and essential BTB domain transcription factors *bric-a-brac 1* and *2 (bab1 and bab2)* (Figure 1B) (Godt and Laski, 1995), and soon also express the signaling molecule *hedgehog (hh)* (Forbes et al. 1996). During prepupal development, TF cells complete germline stem cell (GSC) niche production by specifying cap cells (CCs) at the TF base (Figure 1A; Forbes et al. 1996; Zhu and Xie, 2003; Song et al. 2007; Fu and Spradling, 2026; review Hsu et al. 2019). While niches are being finished, apical ovarian cells migrate down to define ovarioles, enclosing germ cells, and surrounding them with sheaths (Cohen et al. 2002). Subsequently, beginning at the pupal stage, GSCs begin producing daughters at rates that can vary and thereby determine oocyte production by an ovariole throughout adult life.

*Drosophila* females produce eggs at widely varying rates set by the GSCs to maximally utilize available nutrients, while each ovariole maintains about 8 mature germline cysts and 7 follicles in developmental order. They accomplish this by continuously modulating the rate of GSC division, cyst production and follicle progression under the control of insulin and ecdysone signaling and by enforcing checkpoints at the germarium region 2a/2b boundary and at follicle stage 8 (Buszczak et al. 1999; Carney and Bender, 2000; Drummond-Barbosa and Spradling, 2001). After stage 8, the oocyte takes up yolk and lipoprotein particles directly from the hemolymph (DiMario and Mahowald, 1987). The ecdysone that activates post-checkpoint oogenesis is synthesized follicle-autonomously by its ovarian follicle cells, leading to vitellogenesis, eggshell production and ovulation (Terashima et al. 2005; Sun et al. 2008; Ables and Drummond-Barbosa, 2010; König et al. 2011; Morris and Spradling, 2012; Domanitskaya et al. 2014; Knapp et al. 2020; reviewed in Berg et al. 2024).

A germline organelle, the fusome, contains polarized microtubules essential for germline cyst formation in both sexes in *Drosophila*, and participates in apical-basal polarization and oocyte specification (review: Spradling et al. 2022). Recently, similarly polarized germline cysts containing fusome-like structures have been identified in frogs and mammals (Davidian and Spradling, 2025; Pathak and Spradling, 2026). The *Drosophila* fusome contains multiple protein isoforms produced by the the *hu-li tai shao (hts)* gene (Petrella et al, 2007). In GSCs the fusome/spectrosome associates with the cap cells for much of its cell cycle (Figure 1F), before moving to the ring canal (Figure 1G), and fusing with the newly formed fusome (Figure 1H) before being asymmetrically distributed to the daughter cystoblast (Figure 1I). Cysts maintain contact with a series of stationary escort cells in region 2a (Morris and Spradling, 2011) until they reach the start of region 2b halfway through the germarium. Here cysts shed EC membranes and pass two follicle stem cells (FSCs) that each donate a single daughter cell that proliferate to cover their surface (Figure 1A, green).

Rapid germline growth, cyst production, and yolk formation require abundant phospholipids, steroids and other lipids (DiMario and Mahowald, 1987; Sieber and Spradling, 2015; Sieber et al. 2016). In many systems, reserves for these purposes are sequestered in lipid droplets (LDs) (review: Welte, 2015). These dynamic organelles also buffer metabolic stress and protect cells from oxidative damage (Olzmann and Carvalho, 2019; Walther and Farese, 2012; Welte and Gould, 2017). Intestinal enterocytes and adipocytes release lipoprotein particles that circulate in the insect hemolymph bound to lipophorin (Obniski et al. 2018; Carrera et al. 2025). These lipoproteins are taken up by ovarian cells via lipophorin receptors (LpRs), including LpR2, which functions analogously to mammalian LDL receptors to mediate lipoprotein endocytosis (Palm et al., 2012; Sieber and Spradling,, 2015; Matsuo et al. 2019).

Once internalized, lipids are distributed and recycled through endosomal and secretory pathways. Vesicle-mediated transport is central to lipid trafficking, and the exocyst complex is a key regulator of polarized secretion and membrane delivery (Heider and Munson, 2012). Organic anion transporting polypeptides (OATPs) import a variety of neutral lipids into cells including, prostaglandins and ecosinoids (Hagenbuch and Stieger. 2013; Falah et al. 2024). The *Drosophila* genome encodes eight OATP transporters, including Oatp74D which is required for cellular uptake of ecdysone (Okamoto et al. 2018), and Oatp30B whose mammalian ortholog Slco5a1 is of prognostic significance in serous ovarian cancer (Svoboda et al. 2018), but whose function in normal ovaries is not clear.

Here, we demonstrate that the terminal filament takes up and transports lipophilic molecules into the germarium that maintain lipid droplets within the ovariole. Moreover, we show that Oatp30B residing in the terminal filament and cap cells is important for maintaining normal germ cell differentiation into cysts downstream from the GSC.

## METHODS

### Fly Genetics

All *Drosophila melanogaster* stocks were maintained on standard cornmeal agar media at room temperature unless otherwise specified. The Oregon R strain was used as the wild-type control throughout the study. Enhancer trap lines (PZnnnnn) are from Karpen and Spradling (1992); R11A12 is described in (Jenett et al, 2012; www.janelia.org/gal4-gen1). The GAL4 driver lines *bab1-GAL4*, *c587-GAL4*, and *nanos-GAL4* were obtained from the Bloomington Drosophila Stock Center (BDSC). UAS-RNAi lines were obtained from the Vienna Drosophila Resource Center (VDRC), including *Sec6-RNAi* (VDRC #22077), *Sec15-RNAi* (VDRC #35161), *Sec10-RNAi* (VDRC #318483), *Sec5-RNAi* (VDRC #28873), *Exo70-RNAi* (VDRC #103717), *Exo84-RNAi* (VDRC #108650), *Oatp30B-RNAi* (VDRC #110237), *LpR2* (VDRC #25684) and *ACC-RNAi* (VDRC #8105), *ATPα* (VDRC#12330). All genetic crosses were performed at 25 °C unless otherwise noted. For temporally controlled gene knockdown, the temperature-sensitive Gal80 system (*tub-GAL80^ts^*) was used in combination with *bab1-GAL4*. Crosses were maintained at 18 °C to allow GAL80^ts^ mediated repression of GAL4 activity during development. Adult progeny was then shifted to 29 °C to inactivate GAL80^ts^ and permit GAL4 driven RNAi expression at desired developmental stages.

### Egg Laying Assay

Newly eclosed (0–1 day old) flies were maintained in plastic bottles containing molasses plates overlaid with a layer of wet yeast, replaced daily. For egg laying measurements, five pairs of flies were kept per bottle, and the number of eggs laid was counted every 24 hours, in triplicate.

### Immunostaining

Immunostaining was performed on ovaries dissected from adult *Drosophila melanogaster* females. Dissections were carried out in Grace’s insect medium, followed by fixation in 4% paraformaldehyde (PFA) with 0.01% PBST (PBS with 0.01% Triton X-100) for 30 minutes at room temperature. Fixed ovaries were then blocked in 5% Normal Goat Serum (NGS) for 1 hour and incubated overnight at 4°C with primary antibodies diluted in blocking solution. After extensive washing with PBST, samples were incubated with secondary antibodies (1:200 dilution) for 2 hours at room temperature. DNA was counterstained with DAPI (0.5 μg/mL), and samples were mounted in Vectashield mounting medium. Confocal images were acquired using a Leica STELLARIS 8 DIVE microscope.

The following primary antibodies were used: mouse anti-Hts (1:100; DSHB), chicken anti-α5 (1:200; DSHB), mouse anti-LaminC (1:50; DSHB), rabbit anti-Cytochrome C (1:50; Cell Signaling Technology), chicken anti-GFP (1:200; aveslabs) and rabbit anti-Hedgehog (1:500; kindly provided by Dr. Tom Kornberg).

### Nile Red, BODIPY 493/503 and LipidTOX™ Deep Red Staining

To visualize neutral lipids, tissues were dissected in 1× PBS and fixed in 4% paraformaldehyde for 20 minutes at room temperature. Samples were washed three times with 1× PBS and incubated in a Nile Red solution (1 µg/mL in PBS), LipidTOX™ Deep Red (1:500), or BODIPY 493/503 (1 µg/mL in PBS) for 30 to 45 minutes in the dark. After staining, tissues were washed briefly with PBS, counterstained with DAPI (0.5 µg/mL), and mounted in Vectashield for imaging. Images were acquired using a confocal microscope.

### FM1-43 staining

Ovaries were dissected in ice-cold 1× PBS and incubated in FM1-43FX dye (5 µg/mL in PBS; Thermo Fisher) for 15–20 minutes at room temperature in the dark. After staining, tissues were washed briefly with PBS and fixed in 4% paraformaldehyde for 20 minutes. Following fixation, samples were washed three times with PBS and mounted in Vectashield for imaging.

### DHE and MitoSox Staining

To assess reactive oxygen species (ROS) levels, freshly dissected ovaries were incubated in Schneider’s Drosophila medium containing either dihydroethidium (DHE; 5µM, Invitrogen) or MitoSOX™ Red (5 µM, Invitrogen) for 10–15 minutes at room temperature in the dark. After staining, tissues were washed twice with 1× PBS and fixed in 4% paraformaldehyde for 20 minutes. Samples were then rinsed, mounted in Vectashield with DAPI, and imaged using a confocal microscope.

### Quantification of lipid droplet number and volume in the germarium

Lipid droplets in the germarium were quantified using ImageJ. Confocal Z-stacks of BODIPY 493/503, Nile Red and LipidTOX™ stained ovaries were acquired with identical imaging settings across all genotypes. Maximum intensity projections were generated and converted to 8-bit images. The germarium was manually outlined and analysis was restricted to this region.

Images were thresholded using the adjust threshold function with identical threshold parameters applied to all samples within an experiment, and converted to binary masks. Lipid droplets were quantified using the analyze particles function with a size threshold of 0.07 µm² to infinity (corresponding to droplets ≥0.3 µm in diameter). Total LD number per germarium was recorded and data were exported for statistical analysis. Individual lipid droplet volumes were measured using the volume calculator plugin in ImageJ, which estimates volume from the droplet area and optical section thickness, and total lipid content per germarium was calculated by summing all droplet volumes.

### Confocal imaging, image acquisition and processing

A Zeiss LSM 780 laser-scanning confocal microscope (Carl Zeiss) with a 20x and 43x oil-immersion objective (Carl Zeiss) was used for data collection. Confocal z stacks were obtained in 1 μm intervals at a resolution of 1024×1024. Images were processed with Zen software (Zeiss) to obtain maximum projections. Fiji were used for image rotation and cropping.

### Fluorescence Intensity Quantification

Images were acquired using a confocal microscope under identical acquisition settings (laser power, gain, offset, and exposure time) for all samples within an experiment. Imaging parameters were initially optimized using control samples to ensure adequate signal detection without pixel saturation and were then kept constant across all genotypes and conditions. Z-stacks spanning the entire germarium were collected using the same step size for all samples, and maximum intensity projections were generated for analysis. Fluorescence intensity of Hh, DHE and MitoSox staining was quantified using ImageJ. Regions of interest (ROIs) were manually defined over the germarium using identical criteria across samples. Mean fluorescence intensity within each ROI was measured from maximum intensity projections. Background fluorescence was measured from an adjacent region lacking specific signal and subtracted from each measurement. All measurements were performed using identical analysis parameters across samples. Intensity profile graphs were generated using Image J and GraphPad Prism software.

### Statistical Analysis

Graphs and statistical analyses were performed using GraphPad Prism (GraphPad Software). For comparisons between two groups, a two-tailed unpaired Student’s *t*-test was used. A p-value < 0.05 was considered statistically significant. The sample size (n) and statistical test used, are indicated in the corresponding figure legends. Data are presented as mean ± SD unless otherwise stated.

## RESULTS

### The terminal filament and early germarium contain lipid droplets (LDs) and other vesicles

Several recent observations, suggested that the terminal filament and cap cells contain vesicles that have been little studied. Terminal filament cells in ovaries from pupae and newly eclosed adults (Figure 1C) contain abundant small vesicles that stain with Hts, resembling small pieces of fusome (Fu and Spradling, 2026). By 3 days after eclosion, these particles are gone (Figure 1D; Figure S1D). Electron micrographs of TF and cap cell (CC) cytoplasm showed the presence of small vesicles (Figure 1E, arrow). CC membranes contact anterior escort cells (EC), and together these cells comprise the walls of the GSC niche, providing a route for vesicular material to enter downstream germ and somatic cells.

We looked for lipid droplets in these germarium cells like those known to be present in follicle cells and older follicles by staining ovaries with BODIPY 493/503, Nile red or LipidTox which stain primarily neutral lipids (Figure1J; Figure S1B,C). Large LDs were labeled in GSCs, and downstream cells, where they were most abundant in germ cells, and stained preferentially with BODIPY (Figure 1K).

Terminal filament and cap cells transport lipid-rich vesicles in germ cells via exocyst-dependent trafficking:

The stacked organization of the TF suggests that TF cells transfer lipids taken up from the hemolymph into cap cells by undergoing a series of exocytosis and endocytosis events that resulted in the directional movement of cargos (Figure 2A). Membrane fusion events required for these processes depend on the exocyst, a conserved multiprotein complex. When exosomes are released into extracellular space through exocytosis, the exocyst complex represents a key effector of polarized movement, as it is responsible for tethering secretory vesicles to specific plasma membrane domains prior to fusion (Meek et al. 2024). The exocyst is used by most or all cells, and it is critical for cytokinesis in mitotically cycling cells. Adult TF and CC cells do not cycle or divide and appear fully differentiated after the late pupal stage of ovary development. Sec6 is a key exocyst component due to its binding to Sec9 at the plasma membrane and to Rtnl1 in the ER, and for its role in exocyst assembly (Morgera et al. 2012, Dubuke et al. 2015).

**Figure 2:**
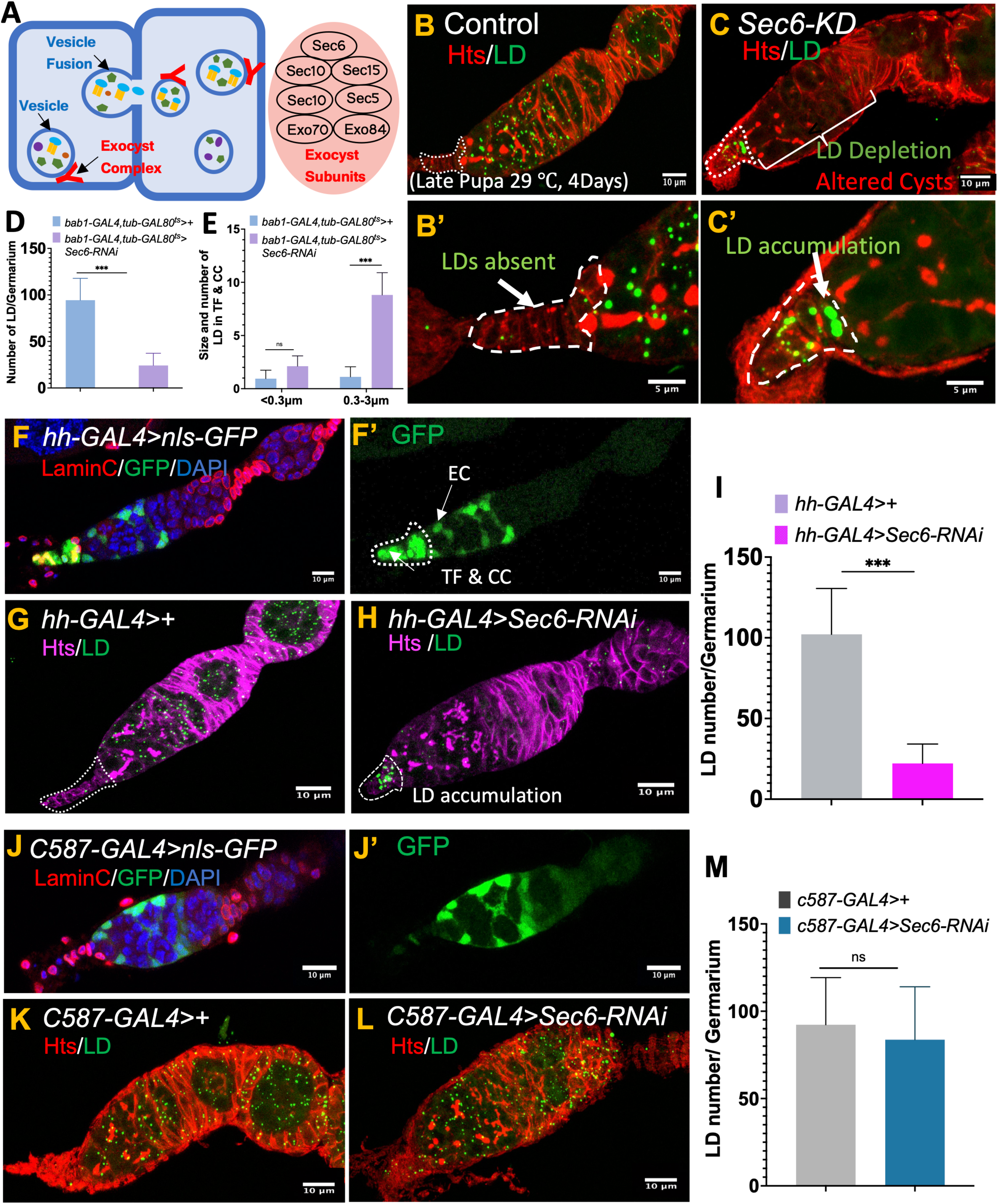
Terminal filament delivers lipids to germ cells through exocyst-dependent vesicle trafficking. **(A)** Schematic diagram of exocytosis showing exocyst complex-mediated cargo delivery. The exocyst complex (marked in red) binds cargo-containing vesicles (blue circle) and tethers them to the plasma membrane, facilitating vesicle fusion and release of cargo to neighboring cells. Exocyst subunits are highlighted in red. **(B)** BODIPY 493/503 staining of control (*bab1-GAL4,tub-GAL80^ts^>+*) germarium shows normal LD distribution at 4d post-eclosion. **(B’)** Higher magnification image of the TF and CC cells (dotted line) in control germarium, showing absence of LDs. **(C)** Conditional knockdown of *Sec6* in terminal filament cells (late pupa-day 3) using the *bab1-GAL4, tub-Gal80^ts^* (Sec6_KD) shows strong depletion of lipid droplets and cyst defects shown via altered fusome, by day 4 post-eclosion. **(C’)** Higher magnification of Sec6_KD germarium (C), showing accumulation of enlarged LDs in TF and cap cells, suggesting impaired lipid export due to disrupted exocytosis. **(D)** Quantification of LD number in control vs. *Sec6_KD* germarium, showing a significant decrease in LD number. Data are shown as mean ± SD; n = 52 germarium examined. **(E)** Quantitative analysis reveals a significant increase in LD number and size within the TF following *Sec6* knockdown, data are shown as mean ± SD; n = 52 germarium examined. **(F-F’)** *hh-GAL4* driven *nls:GFP* marks TF, CC and EC. (**G-H**) *hh-GAL4* driven *Sec6* KD **(H)** leads to accumulation of LDs only in TF and CC cells, not in escort cells, with loss of LDs from the germarium, compared to control **(G)**. (**I**) Graph represents significant reduction of LDs from *hh-GAL4* driven *Sec6 KD* germarium, data are shown as mean ± SD; n = 50 germarium examined. (**J, J’**) *c587-GAL4* driven GFP marks escort cells in germarium. (**K-L**) *Sec6* knockdown **(L)** in escort cells driven by *c587-GAL4* disrupts cyst and follicle formation, but LD levels in germ cells remain largely unaffected, compared to control (**K**). **(M)** Quantification of LDs signal in germ cells shows no significant difference between control and escort cell specific *Sec6* knockdown, supporting a limited role for escort cells in lipid transport, n=52 represents number of germarium analyzed with mean ± SD. Statistical significance was determined using unpaired two-tailed t-test, (ns = not significant, *P < 0.05, **P < 0.01, ***P < 0.001, ****P < 0.0001).

Consequently, we knocked down the gene encoding Sec6 by using the *bab1-GAL4* driver specific for TF and cap cells (Figure S1A) to drive anti-*Sec6* RNAi, and GAL80^ts^ repressor to provide temporal control. Late pupae were shifted to 29 °C to inactivate GAL80^ts^, and both controls and KD animals were analyzed after 4 days at 29 °C using BODIPY 493/503 staining. LD levels remained high in controls (Figure 2B), but were strongly reduced throughout the germarium and follicles in KD ovarioles (Figure 2C,D). Moreover, while LDs were depleted from GSCs and downstream germ cells, large LDs accumulated specifically in posterior TF and cap cells in *Sec6* knockdown animals where they are rarely observed in controls (Figure 2C’ vs 2B’). The size increase was confirmed by counts based on LD size (Figure 2E). LD accumulation is predicted by the transcytosis model of TF import.

Extending the study of *Sec6* KD to ten days showed that these animals strongly reduced egg production (Figure S2A). To verify that changes in vesicle trafficking were involved in these effects, we used FM1-43 dye which labels vesicles undergoing membrane recycling and regulated exocytosis (Ramachandran and Budnik, 2010). FM1-43 has been previously employed in studies of synaptic activity and neurogenesis, where it marks vesicle turnover during neurotransmitter release. TF and CC cells both showed staining indicative of active exocytosis (Figure S2B). However, *Sec6* knockdown ovarioles (Figure S2C) had a significantly reduced FM1-43 signal in TF cells (Figure S2F). A strong reduction of LDs in *Sec6* KD ovarioles similar to that observed with BODIPY staining was also documented when the less selective lipid dye Nile Red was used (Figure S2, D,D’,G). LDs again accumulated in the TF (Figure S2E-E’,G).

Knockdown of 5 other exocyst components (*Sec10*, *Sec15*, *Sec5*, *Exo70*, *Exo84*) also resulted in LD depletion (Figure S3 A-F, G), but only Sec6 depletion consistently caused LD backup near the CC/GSC border. *Sec6* knockdown also abrogated normal cyst and follicle development (Figure S2H-I). Together these finding show that exocyst-mediated lipid delivery from TF and CC is essential for germ cell and cyst development. In the absence of transport, ongoing metabolic demands of growing germ cells apparently leads to the exhaustion of their lipid reserves stored in LDs, causing them to disappear.

### Escort cells contribute little to supporting LDs in the germarium

To investigate whether the use of the *bab1-GAL4* driver was needed for the observed changes in LDs in TF, CC and downstream germarium, we used a *hh-GAL4* driver, which is also expressed in TF, CC, as well as in escort cells (Figure 2F-F’). Knockdown of *Sec6* using *hh-GAL4* resulted in a pronounced depletion of lipids from the germarium (Figure 2G-I).

Furthermore, lipid droplets accumulated specifically in posterior TF and in CC cells, but not in escort cells (Figure 2G-H). This provided further reinforcing evidence that TF and cap cells, but not escort cells, are important for lipid entry based on endocytic activity. We also investigated the contribution of germarium somatic cells using the *c587-GAL4* driver to selectively inactivate *Sec6* in ECs, FSCs and early follicle cells, but not TF and cap cells (Figure 2J, J’). No loss of LDs or buildup of LDs in any germarium cells were observed (Figure 2K-M).

### Kinetics of LD loss and TF accumulation

We investigated the kinetics of LD loss following *Sec6* inactivation using *bab1-GAL4: GAL80^ts^*- mediated RNAi expression. Figures 3A-J shows images of a representative germarium imaged every 6 hours after shifting the temperature to 29 °C. The left hand axis shows that total LD volume (in droplets of measurable size) is initially negligible in TF and CC cells, but has increased after 6 hours, and begins to plateau after 12 hours, reaching a final value of about 0.6 μm^3^ within the estimated 15 total TF and CC cells. The right hand axis shows that the total volume in LDs, 4.4 μm,^3^ began to decline between 12 and 18 hours in the approximately 100 germ cells in a germarium. The reciprocal nature of these curves provides further assurance that the TF and CC cells normally import lipophilic materials needed to maintain the large germ cell LDs.

**Figure 3:**
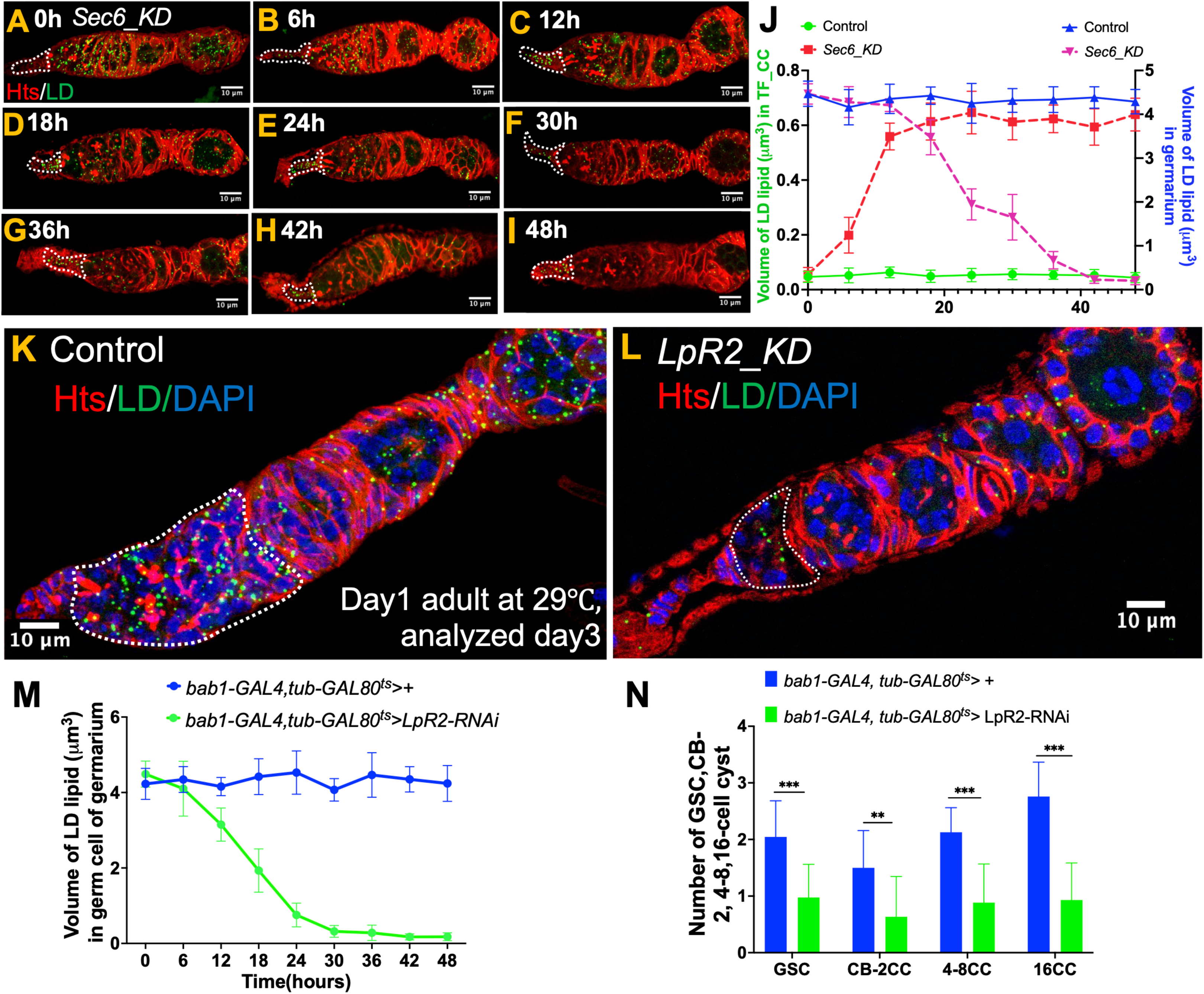
Kinetics of Sec6 and LpR2 dependent lipid transfer from terminal filament to the germarium. **(A-I)** Hts and LD staining (BODIPY 493/503) of germarium at 6-hour interval from 0 to 48 hours post-eclosion following *Sec6* knockdown using *bab1-GAL4, tub-GAL80^ts^* (*Sec6_KD*), with temperature shift to 29 °C. LDs accumulate within the TF from 12 hours, while their levels are markedly reduced in germ cells starting from 18hours. **(J)** Quantification shows volume of lipid in LDs of germarium (right Y-axis) starts to decline from ∼18 hour onwards and by 30-36 hours showed complete depletion compared to control. Quantification of lipid volume in TF cells (left Yaxis) shows a steady increase from 12 hours to 48 hours in *Sec6* knockdown compared with control. Data are presented as mean ± SD, n = 52 germarium from each genotype. **(K-L)** *LpR2_KD* knockdown (*bab1-GAL4,tub-GAL80^ts^>LpR2-RNAi*) in terminal filament and cap cells on adult day1 and analyzed on day3, leads to a reduction in lipid droplet number within the germarium, a smaller region 1 and 2A with less-branched fusomes (dashed outline), and defects in cyst development. **(M)** Graph showing the reduction in lipid droplet volume within germ cells over time (0–48 h) following *LpR2* knockdown in terminal filament and cap cells, lipid reduction begins around 12 h post-knockdown, data shows mean ± SD, n = 48 shows number of germarium analyzed. **(N)** Graph represents quantification for GSC, CB-2CC, 4-8CC and 16CC cyst defects in *LpR2_KD* compared to control, data are shown as mean ± SD, n=44 number of germarium examined. Statistical significance was determined using unpaired two-tailed t-test, (ns = not significant, *P < 0.05, **P < 0.01, ***P < 0.001, ****P < 0.0001).

The number of LDs in both the germarium, TF and CC cells was also determined at each time point (Figure S4A-B). These values changed in parallel with volume. Regionally, however, the rate of LD declines were not identical within the four main germarium "regions": region 1, 2a, 2b, and 3 (Figure S4C). Finally, the number of 16-cell cysts in region 2b, was counted based on fusome staining (Figure S4D; Figure 4H). The number of these cysts remained little changed until 30 hours after which it fell almost to baseline in the next 18 hours.

**Figure 4:**
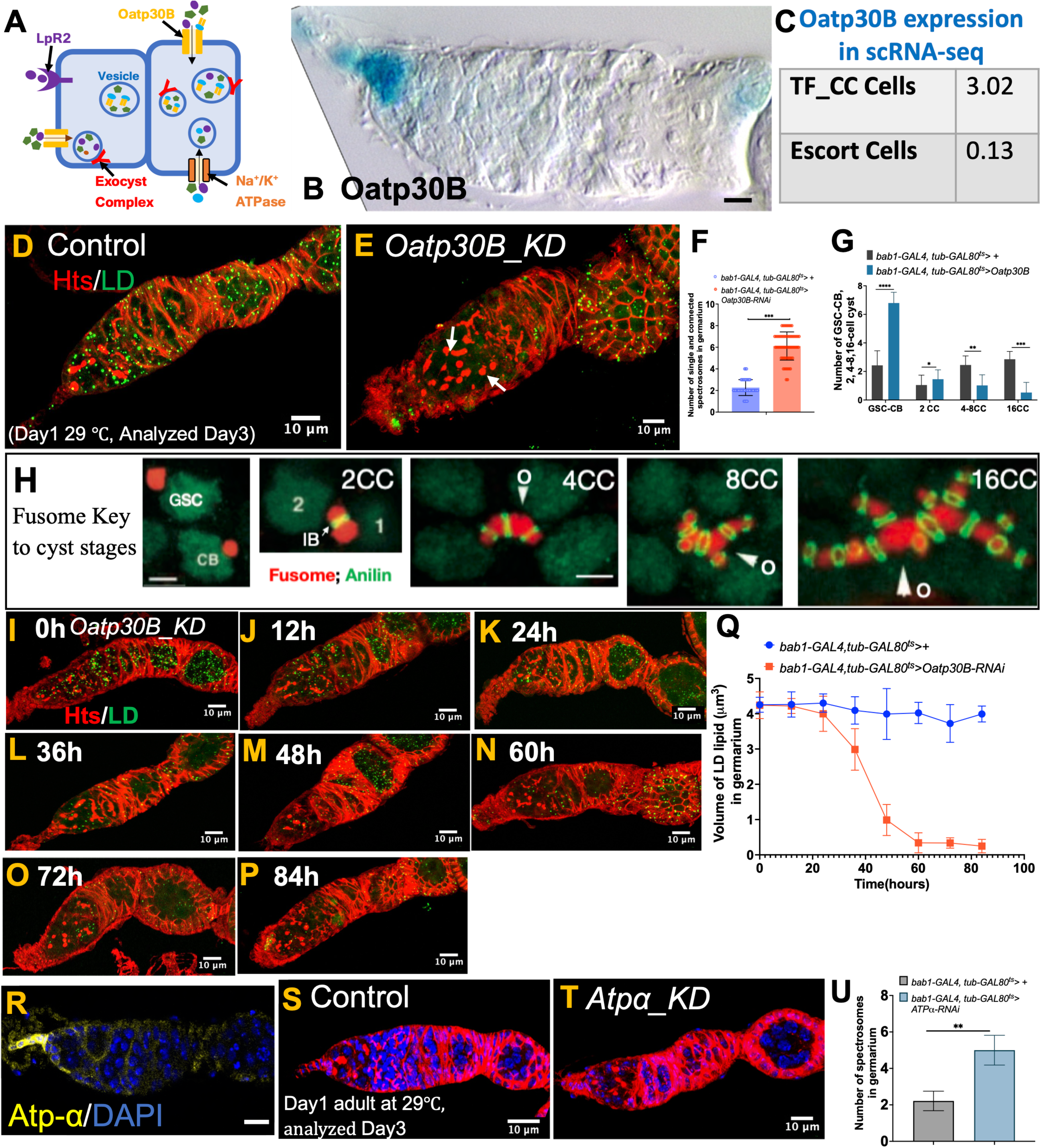
Oatp30B mediated transport in the terminal filament regulates cyst development. **(A)** Schematic representation of Oatp30B, a membrane-bound transporter, may facilitates extracellular cargo uptake, which can be subsequently packaged into intracellular vesicles. These cargo-packed vesicles are tethered to the plasma membrane through the exocyst complex, driving vesicle fusion and exocytotic release of cargo to neighboring cells. LpR2 (lipoprotein receptor) and Na /K -ATPase (membrane ion pump) are depicted as additional plasma membrane associated components. **(B)** TF and germarium from enhancer trap line Oatp30B (PZ00078) reveals strong enrichment in terminal filament cells. **(C)** scRNA-seq data shows high expression of Oatp30B in TF and cap cells as compared to escort cells of the germarium. **(D)** LDs labeled with BODIPY 493/503 and Hts in control germarium. **(E)** Knockdown of Organic anion transporting polypeptide 30B (*Oatp30B*) in TF and CC was achieved using *bab1-GAL4,tub-GAL80^ts^* with RNAi induced at 29 °C from day 1 and analyzed on day 3. LDs were strongly reduced, moreover, knockdown caused increase in germ cells containing spectrosomes. **(F)** Quantification showing increase in number of single and connected spectrosomes till region2A in *Oatp30B_KD* compared to control, n=58 number of germarium analyzed, mean ± SD. **(G)** Quantification of germline cyst progression defects in *Oatp30B_KD,* including GSC-CB, 2-cell cysts, 4–8-cell cysts, and 16-cell cysts. *Oatp30B_KD* exhibits a significant accumulation of GSC-CB with a concomitant reduction in 16-cell cysts, indicative of impaired cyst differentiation, data are shown as mean ± SD from n=58 germarium analyzed. **(H)** Representative images depicting fusome morphology at successive stages of germline cyst development, from the undifferentiated germline stem cell (GSC) and cystoblast (CB) through 2-cell (2CC), 4-cell (4CC), 8-cell (8CC), and 16-cell (16CC) cyst stages, illustrating the progressive branching and growth of the fusome during cyst formation. images from de Cuevas and Spradling, 1998. **(I-P)** Oatp30B was knocked down using *bab-GAL4, tub-GAL80^ts^* starting on day 1 of adult eclosion, and lipid droplets were assessed at 12-hour intervals. A noticeable reduction in lipid droplet number was observed from 24 hours onwards, with a gradual decrease over time. **(Q)** Graph showing reduced lipid droplet volume in the germarium from 24 hour onwards following *Oatp30B* knockdown compared to control. Data are shown as mean ± SD from n=58 germarium. **(R)** Strong expression of Na /K -ATPase stained with Atp-α in terminal filament cells. **(S-T)** *Atp*_α_ knockdown using the same driver as for *Oatp30B*, with RNAi induced at 29 °C from day 1 and analyzed on day 3, shows increased spectrosome extending through region 2A of the germarium compared to control. **(U)** Quantification of spectrosome number per germarium in Atpα_KD compared to control, revealing a marked increase in spectrosome-containing cells, indicative of a differentiation block at the GSC or cystoblast stage, data are shown as mean ± SD from n=48 germarium. Statistical significance was determined using an unpaired two-tailed t-test (ns = not significant, *P < 0.05, **P < 0.01, ***P < 0.001).

### LpR2 is needed to maintain LD levels and germ cell development

The major low-density lipoprotein (LDL) receptor LpR2 (Matsuo et al., 2019) functions late in oogenesis during lipid yolk formation (Parra-Peralbo and Culi, 2011; Sieber and Spradling, 2015). We investigated if LpR2, which is expressed by RNAseq in TF and cap cells at low levels (Sladina et al. 2021), plays a role in TF-mediated import. When LpR2 was knocked down in the TF using the *bab1-Gal4* driver, and *GAL80^ts^* repressor, LD levels were reduced to low levels by day 2 (Figure 3K,L). A complete time course (Figure S5A-H) showed that LD number was reduced to a basal level by 30 h (Figure S5J). We also measured total germ cell LD volume over time to estimate the rate of LD-mediated lipid transfer from terminal filament to the germarium (Figure 3M). LD lipid volume declined by about 1.6 µm³ between 6 and 12 hours post-knockdown, followed by a further reduction of ∼1.4 µm³ during the subsequent 6 hour interval.

The above measurements show a steady decline in LD volume that probably results from a cut off of precursor supplies for LD maintenance from the TF. However, the developmental affects of losing LpR2 preferentially in the TF and CC cells, are different from the those of losing Sec6. Loss of LpR2 in the TF apparently caused some cysts to revert back toward GSCs, as indicated by the appearance of GSC-like single germ cells in region 2a (Figure S5A-C). By 24h cysts were reduced in number (Figure S5E) and their fusomes were less branched compared to controls (Figure S5F-H, arrows). By 30h, LDs were mostly gone (Figure S5J) and cyst development was sparse and abnormal throughout the germarium (Figure 3N, Figure S5F-H).

Thus, TF-specific depletion of LpR2 may not only disrupt the delivery of LD precursors, but differentially affect the levels of one or more molecules required for cyst production and development, compared to sec6 depletion.

### Oatp30B is essential for cyst differentiation, LD acquisition and Hh signaling

The terminal filament and cap cells preferentially express Oatp30B (Figure 4A,B) as confirmed by scRNAseq (Figure 4C, Sladina et al 2021). We knocked down this gene using *bab1-GAL4* and GAL80^ts^ beginning in newly eclosed females. Over the next 3 days, normal cyst production in germarium region 2a gradually deteriorated (Figure 4D-E, I-P). There was a least a 3-fold increase in the number of single cells and 2-cell cysts (Figure 4F) and a decrease in the number of 8 and 16-cell cysts in region 2a (Figure 4G) as determined by fusome morphology used to identify cyst stages (Figure 4H). LDs were lost between 24 and 48 hours after temperature shift (Figure 2I-P), a rate of loss of about 2.4 µm³/24 hours (Figure 4Q). Thus, Oatp30B imports molecules into the TF which are necessary for ongoing TF maintenance of LDs, as well as a unique molecule(s) essential to maintain germ cell development. Oatp30B KD also reduced the level of Hh in the TF (Figure S6 B-C, D). However, Hh alterations cannot explain the drastic inhibition of germ cell differentiation, as Hh produces a very different effect following disruption of *Oatp30B^KD^* (Forbes et al. 1996a).

Some Oatp family members specifically interact with the Na-K-ATPase (ATPα) in transporting cargos (Torrie et al. 2004; Reinhard et al. 2011). Antibody staining revealed that ATPα protein is highly enriched in the terminal filament, and weakly expressed on the cell membranes at the outer surface of developing region 2a cysts (Figure 4R). Knocking down *ATPα* in the TF using *bab1-GAL4 tubGAL80^ts^* on day 1 had strong effects by day 3 (Figure 4S,T). The total number of cysts in region 2a including 16-cell cysts was reduced while early cysts were increased (Figure 2U). While it is known that loss of membrane potential can activate Hh signaling (Emmons-Bell and Hariharan, 2021), Hh overexpression (Forbes et al. 1996a) could not have caused the observed cyst reduction.

### Lipid droplets in the germarium protect against mitochondrial ROS accumulation

We hypothesized that the continuous lipid supply from the terminal filament is required not only to meet energetic demands of early germ cells, but serve functions beyond energy storage. We have observed that LDs in early germ cells are frequently in close proximity to mitochondria in the germarium (Figure 5A, encircled in yellow). This spatial association is consistent with earlier reports suggesting that LD-mitochondria contacts facilitate fatty acid transfer, enabling mitochondrial respiration and regulating oxidative homeostasis (Rambold et al., 2015; Bailey et al. 2015; Nguyen et al., 2017; Avila et al. 2025). Disruption of these contacts or LD content has been linked to elevated mitochondrial reactive oxygen species (ROS) and organelle dysfunction.

**Figure 5:**
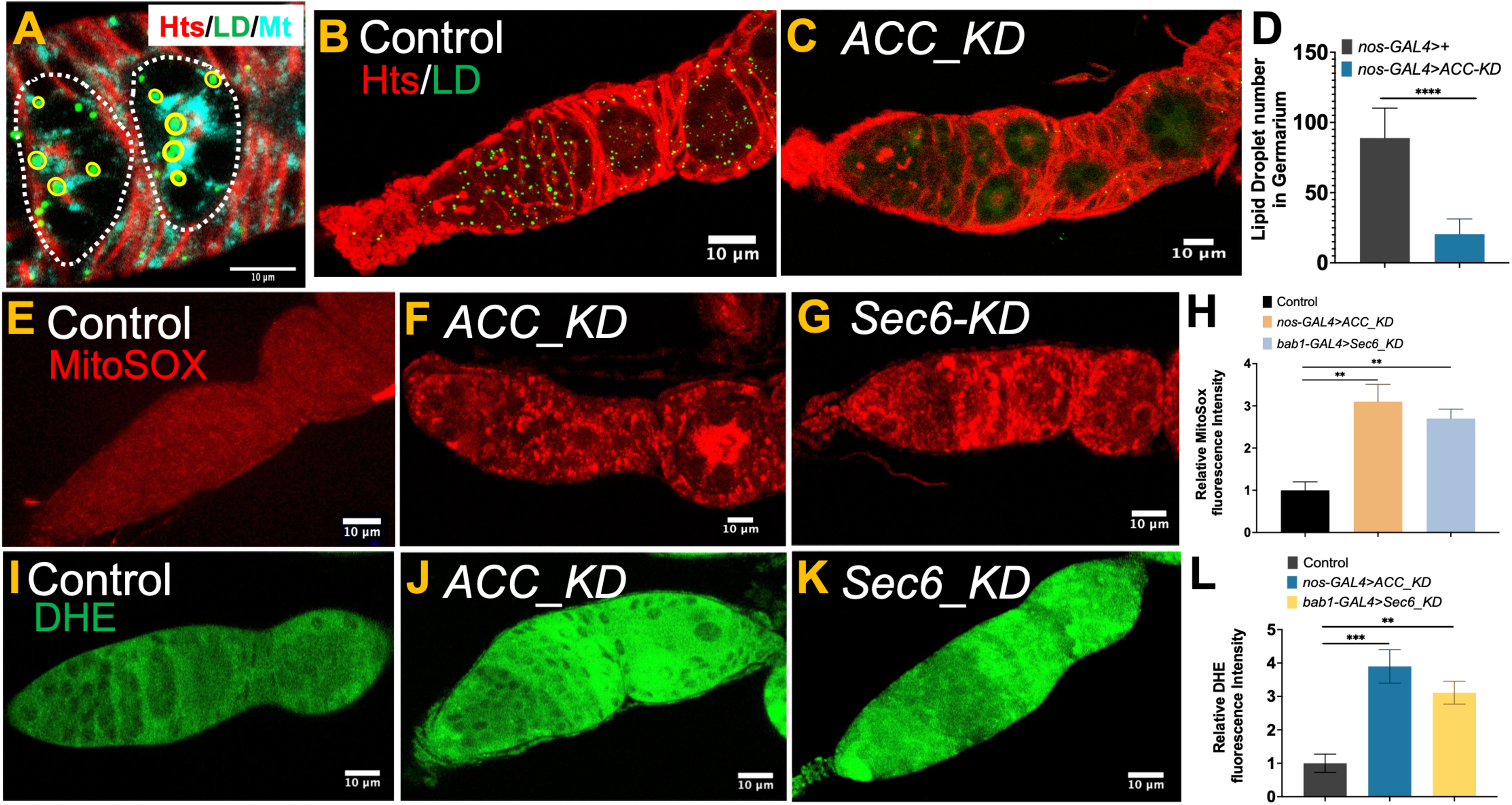
Lipid droplets protect early germ cells from mitochondrial ROS accumulation. **(A)** Control germarium showing close association of lipid droplets (BODIPY 493/503) with mitochondria (Cyto C). **(B–C)** Representative images of control (B) and germ cell-specific ACC knockdown (C) germarium driven by *nanos-GAL4*, showing severe loss of lipid droplets upon depletion of the lipogenic enzyme Acetyl-CoA Carboxylase (ACC). **(D)** Quantification of LD levels in germ cells shows a significant reduction following *ACC* knockdown. Data are presented as mean ± SD, n =42 individuals examined.**(E–G)** MitoSOX staining reveals elevated mitochondrial ROS in *nanos-GAL4* driven *ACC* knockdown (F) and TF-specific *Sec6* knockdown (G), compared to control (E). **(H)** Quantification of MitoSOX signal intensity shows significant ROS increase in both *ACC* and *Sec6*-depleted germarium. Data are shown as mean ± SD, n=48 germarium analyzed from each genotype. **(I-K)** DHE staining detects increased ROS levels in *nanos-GAL4* driven *ACC* knockdown (J) and TF-*Sec6* knockdown (K) germaria relative to control (I). **(L)** Quantification of DHE signal intensity confirms significantly elevated ROS in both experimental conditions, supporting a protective role for LDs against oxidative stress. Data are shown as mean ± SD from n = 48 analyzed germarium. Statistical comparisons were performed using unpaired two-tailed Student’s t-test. ns = not significant, *P < 0.05, **P < 0.01, ***P < 0.001

To directly test whether loss of lipid droplets leads to ROS accumulation, we knocked down the lipogenic gene Acetyl-CoA Carboxylase (ACC) in germ cells using *nanos-GAL4*. ACC is a key enzyme in de novo fatty acid synthesis and is essential for neutral lipid production. ACC knockdown resulted in a dramatic reduction or complete loss of LDs throughout the germarium (Figure 5C vs 5B,D). We next assessed mitochondrial ROS using MitoSOX and DHE staining. Both assays showed significantly elevated ROS levels in *ACC*-depleted germarium compared to controls (Figure 5E,F,H; 5I-J,L), indicating oxidative stress resulting from lipid depletion.

To determine whether this ROS phenotype also arises when LD transport from the TF is impaired, we examined TF-specific knockdown of *Sec6*. As previously shown, *Sec6* knockdown blocks vesicular transport of lipids to the germarium. Consistent with our hypothesis, *Sec6*-RNAi in TFs also led to elevated mitochondrial ROS in germ cells, as measured by both MitoSOX and DHE (Figure 5E,G,H;5I,K,L), which ultimately affects cyst development and follicle formation.

## DISCUSSION

### The terminal filament imports lipophilic molecules

Our experiments demonstrate that a major function of the *Drosophila* terminal filament and cap cells is to import lipophilic molecules necessary to maintain germ cell lipid droplets and regulate normal cyst formation within the ovariole. Molecules entering via the TF are delivered to the GSC, downstream germ cells, and anterior escort cells (Figure 6A). The terminal filaments allow ovarioles to operate semi-autonomously, almost as independent small ovaries (Figure 6B). This promotes the ovarioles autonomy, so that a problem in one ovariole that slows follicle movement does not impede oogenesis more widely. This parallelization of gametogenesis is similar to the role of seminiferous tubules in the testis (Yoshida, 2020), and is found in many animals.

**Figure 6:**
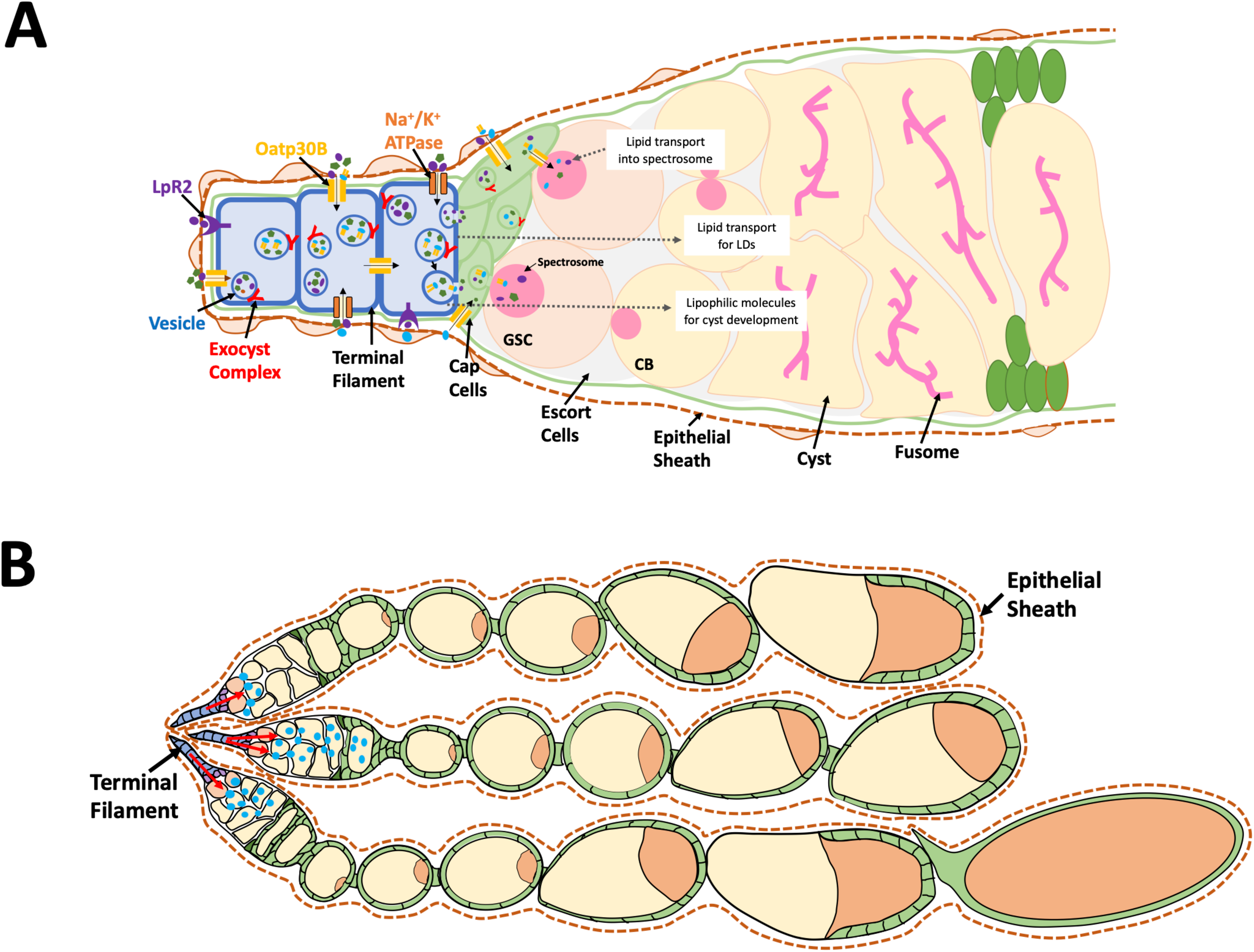
Model for terminal filament uptake, intercellular transport and delivery of cargos to germarium. **(A)** Terminal filament (TF) cells actively incorporate molecules from the hemolymph via transporter (Oatp30B) and receptor (LpR2), process and package them, and deliver the resulting cargo including lipids, and organic solutes to ovariolar germ cells via vesicle-mediated trafficking through transcytosis. **(B)** Terminal filament mediated transport of lipophilic molecules (shown in blue) regulates follicle development in an ovariole-specific manner within the epithelial sheath.

Ovarioles maintain similar patterns of follicle development without compromising their independence. Cysts and follicles maintain a strict sequential order increasing from the anterior GSCs to mature eggs at the posterior. Each ovariole hosts approximately 8 cysts in the germarium and 7 follicles. But the similar stages between ovarioles are not precisely synchronized and only ovulation shows a full coordination of all 32 ovarioles to ensure passage of one egg at time. Similar, but unsynchronized development depends on checkpoints that operate at the 2a/2b germarium border (Drummond-Barbosa and Spradling, 2001), stage 8 (Buszczak et al. 1999) and at egg maturity to limit excess accumulation (see Berg et al. 2024).

### Terminal filament import is important for germ cell metabolism

A major function of the terminal filament is to import precursors that make it possible to maintain substantial LDs in each germ cell. These precursors likely include triglycerides and cholesterol transported through the hemolymph and taken up by TF cells, as no other source is available. GSC division, cyst formation and growth under rich nutritional conditions would require a substantial input of lipids. This may explain why species that produce an exceptionally large number of offspring and at high rates such as honeybee queens have particularly long and cell-rich terminal filaments (Cullen et al 2023). Additionally, LDs have the potential to buffer excess free fatty acids, distribute lipids to mitochondria in a regulated manner, and prevent lipotoxic stress. TF-mediated import allows the maintenance of large LDs in each germ cell that might help explain why *Drosophila* female GSCs are less dependent on glycolysis than many stem cells (Wessel and Drummond-Barbosa, 2026) and the effects of adipocyte gene expression on female GSC and germ cell behavior (Zike et al. 2026).

We estimated the rate of lipid import based on the rate of LD loss following terminal filament knockdown of sec6, LpR2 or Oatp30B. The loss of about 0.13 μm^3^/hr (Figure 3J,M; Figure 4Q), indicates that the TF provides a substantial ongoing supply of triglycerides and other components needed to maintain the steady state LD level observed under normal circumstances. The lag time before LD depletion began varied between 0 hr (LpR2) and 24 hr (Oatp30B), which might reflect differences in RNAi efficiency or physiological differences.

### The terminal filament supports early follicle development within its ovariole

Our studies allowed us to infer that the TF transports some molecules that act as specialty materials, or regulators, rather than simply prerequisites for growth. The small fusome-like particles found only in pupa and newly eclosed adults may supplement fusomes of early progeny, before this function can be taken over by adult GSCs (Figure 1C-D). Disrupting transport via the TF did more than reduce normal cyst production in region 2a, but caused it to become abnormal. LDs contain a large number of lipid species, including regulators like steroids and prostaglandins as well as peptides and protein (Welte, 2015, Welty and Gould, 2017; Thomalla et al. 2025). The developmental effects in our experiments may reflect gene-specific changes in the relative amounts of multiple molecules that act on germline and somatic cell development, as well as well hormones such as ecdysone (Drummond-Barbosa, 2019) and juvenile hormone (Kurogi et al. 2023; Zhu et al. 2024).

The Oatp30B transporter was of particular interest in this regard due to its high expression in the terminal filament, and strong requirement to maintain cyst formation. Slco5a1, the mammalian ortholog of Oatp30B is expressed in pituitary endocrine cells as well as oocytes and granulosa cells (Human protein atlas, 2026). The the function of neither Slco5a1 or Oatp30B is currently well characterized. Slco5a1 expression in serous ovarian cancer is associated with better outcomes (Svoboda et al. 2018). A previous study reported that Oatp30B is one of three *Drosophila* Oatp family members capable of transporting the plant cardiac glycoside ouabain (Torrie et al. 2004). Our experiments showed that Oatp30B likely transports one or more molecules needed to maintain germline cyst development.

### The spectrosome may shuttle materials from the TF-CC base into the fusome during the GSC cycle

The terminal filament/cap cells have long been known to interact with the GSC’s fusome (spectrosome) during its asymmetric division cycle (deCuevas and Spradling, 1998). During the long G2 phase, the fusome/spectrosome remains adjacent the anterior cap cells (Figure 1F). At M phase it migrates to the ring canal following incomplete cytokinesis and fuses with the plug of new fusome destined mostly for the cystoblast that is located there (Figure 1G-I). We propose that while docked at the TF/CC base during the long G2 phase, the GSC’s spectrosome receives specific molecules transported by the TF (Figure 6A, arrow). These materials may be transferred to the cystoblast daughter’s fusome and prepare it for normal cyst formation.

### Terminal filament-regulated import may be important for rejuvenation and metabolism

A major challenge for meiosis and oogenesis is germ cell rejuvenation (Holliday, 1984; Kirkwood, 1987; Cox and Spradling, 2003; Unal et al. 2011; reviews: Xiao and Unal, 2025; Spradling et al. 2025). Recent work suggests that rejuvenation in mouse oogenesis starts as the PGCs begin cyst formation, much earlier than previously supposed (Pathak and Spradling, 2026). A sequestered route into the GSC and anterior germarium might allow close control of the raw materials destined to construct new oocytes. In budding yeast, the spore membranes are initiated first spindle poles by a microtubule-dependent cap, and begin to grow by attracting vesicles even before acquiring a meiotic nucleus (Neinman, 2011). Animals like *Drosophila* may begin to rebuild membranes even before meiosis, using imported lipids, as well as and the spindle microtubule scaffold, vesicles and biosynthetic capacity present in the ER-rich fusome.

Our experiments showed that another function of imported lipids is to protect germ cells in the germarium, a critical region for rejuvenation, from oxidative damage. Both ACC knockdown in germ cells and TF-specific Sec6 knockdown increased mitochondrial ROS, demonstrating that disruption of LD accumulation compromises redox homeostasis. Lipid provisioning enables biosynthetic and energetic demands while safeguarding germ cells from oxidative damage. LD–mitochondria contacts facilitate fatty acid transfer and protect against ROS accumulation (Nguyen et al., 2017; Rambold et al., 2015). In the absence of LDs, free fatty acids can undergo peroxidation, triggering a cascade of oxidative damage (Listenberger et al., 2003; Bailey et al., 2015). In *Drosophila* and mammalian neurons, lipids play a critical role in suppressing oxidative damage through an interaction involving lipid exchanges with glial cells (Liu et al. 2015; Moulton et al. 2021; review: Goodman et al. 2024). The association of GSCs and early germ cells with escort cells, squamous cells that resemble glial cells, may contribute to their enhanced, LD-dependent oxidative resistance.

## ACKNOWLEDGEMENTS

We thank Mahmud Siddiqi for assistance with light microscopy and Mike Sepanski for electron microscopy. Dianne Williams assisted with *Drosophila* stock management. We are grateful to Asya Davidian, Haolong Zhu, Madulika Pathak, Ashish Tiwari, Yunpeng Fu, Andy Mao and other members of the Spradling laboratory for discussions. Allan Spradling is an Investigator of the Howard Hughes Medical Institute, and HHMI provided support for these studies. The Carnegie Institution for Science has generously supported the Spradling lab in Baltimore for many years, for which we are grateful.

## Notes

### Competing Interest Statement

The authors have declared no competing interest.

### Summary of Updates

The paper was narrowed to focus on the import of lipophilic molecules by the TF. Material related to any roles in ecdysone and cytokines were removed and will be published elsewhere. The writing was revised to reflect this organization. More quantitation of there kinetic data was added along with statistical support.

