## Supplemental figures and legends for "The *Drosophila* ovarian terminal filament imports lipophilic molecules that support cyst and follicle development within the ovariole"

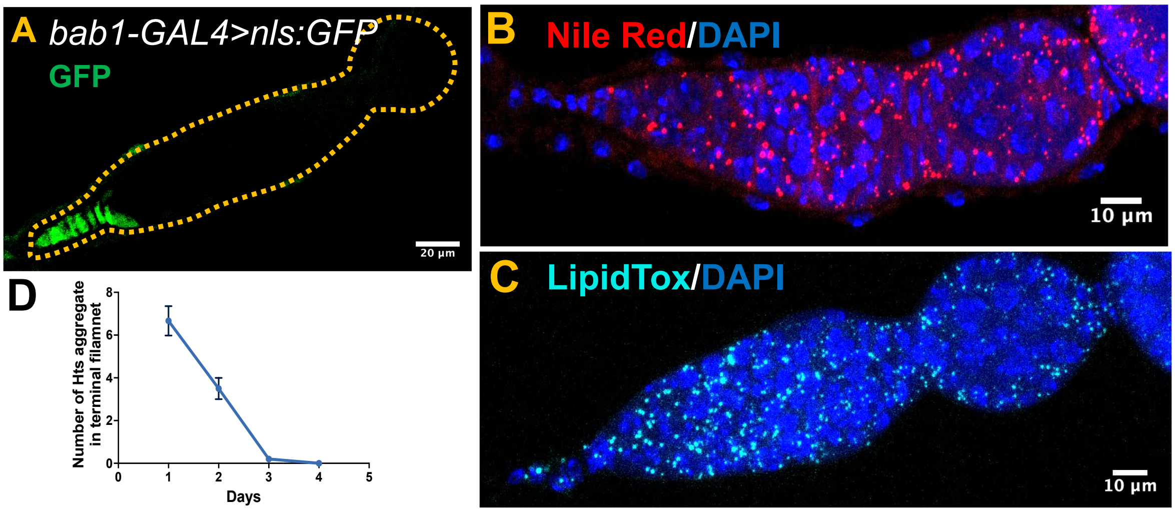


**Figure S1: Localization of lipid droplets and Hts-positive aggregates in the germarium.** **(A)** Germarium tip expressing nls:GFP under *bab1-GAL4*, labeling TF and CC cells. **(B-C)** Germarium stained with Nile Red (B) and LipidTOX (C), highlighting lipid droplet distribution throughout the germarium. **(D)** Decline and disappearance of Hts aggregates in the TF by day3 of adulthood.

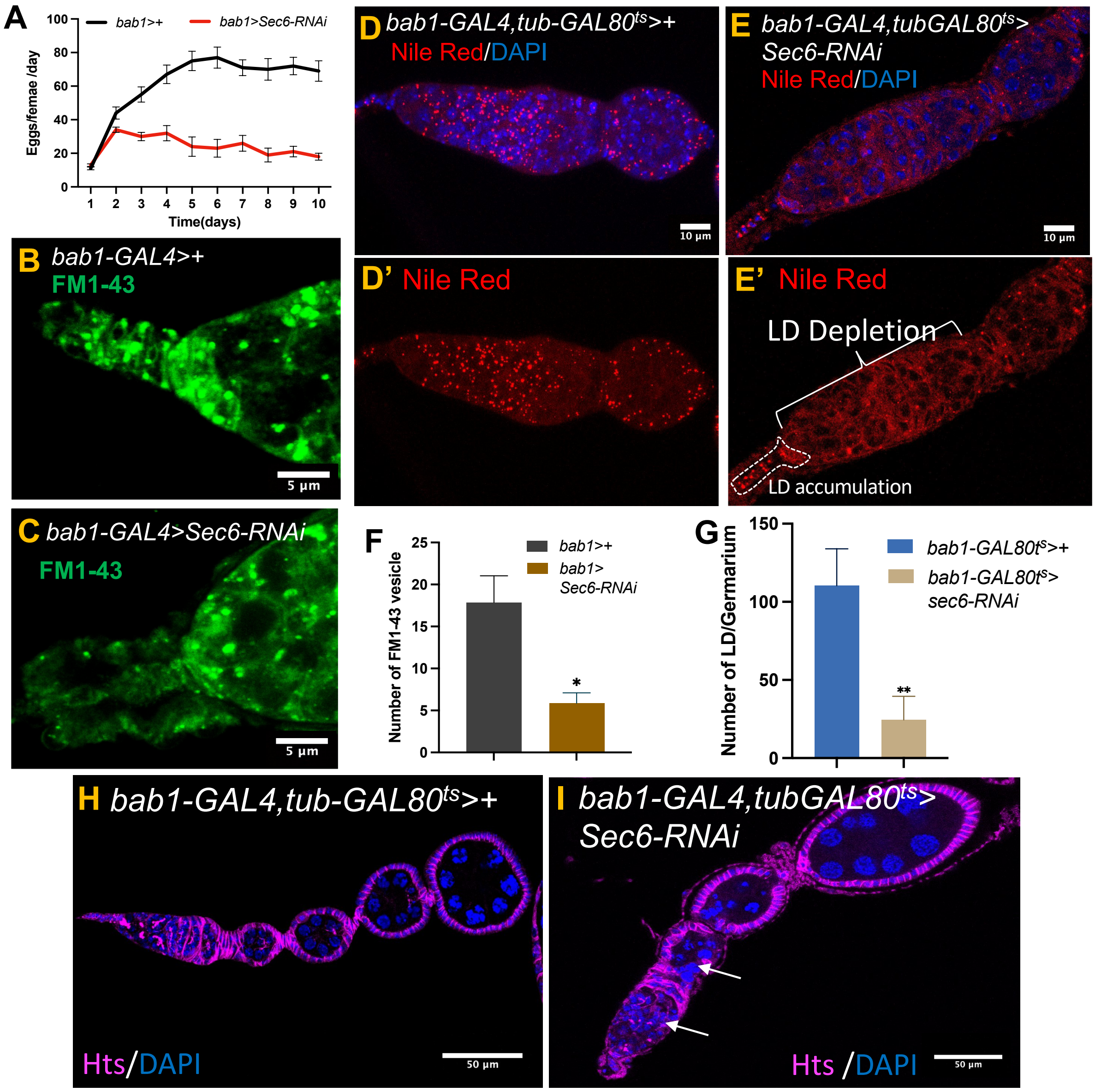


**Figure S2: Active exocytosis in terminal filament and cap cells with lipid droplet accumulation.**

**(A)** Egg laying assay showing a significant reduction in the number of eggs produced per female per day following *Sec6* knockdown with *bab1-GAL4*. **(B)** FM1-43 dye staining in control germarium shows strong signal in TF and cap cells, indicative of active vesicle trafficking and membrane recycling. **(C)** FM1-43 staining is markedly reduced in TF and cap cells following *Sec6* knockdown, suggesting impaired exocytosis. (**D, D'**) LDs stained with nile red in control germarium. (**E, E'**) Depletion of nile red positive LDs in germarium of *Sec6* knockdown in TF and CC, along with accumulation of LDs in TF and CC cells. **(F)** Quantitative analysis of FM1-43 fluorescence intensity confirms a significant reduction in

TF/cap cells in *Sec6* knockdown samples compared to control. Data are presented as mean ± SD, n = 32 germarium from each condition. (**G**) Graph represents reduction of LDs in Sec6 knockdown. Data are presented as mean ± SD, n = 38 germarium from each genotype. **(H)** Control ovariole with normal cyst and developing follicles. (**I**) *Sec6_KD* showing defective cyst and follicle development marked with arrow. Statistical significance was determined using unpaired two-tailed t-test, (ns = not significant, *P < 0.05, **P < 0.01, ***P < 0.001, ****P < 0.0001).

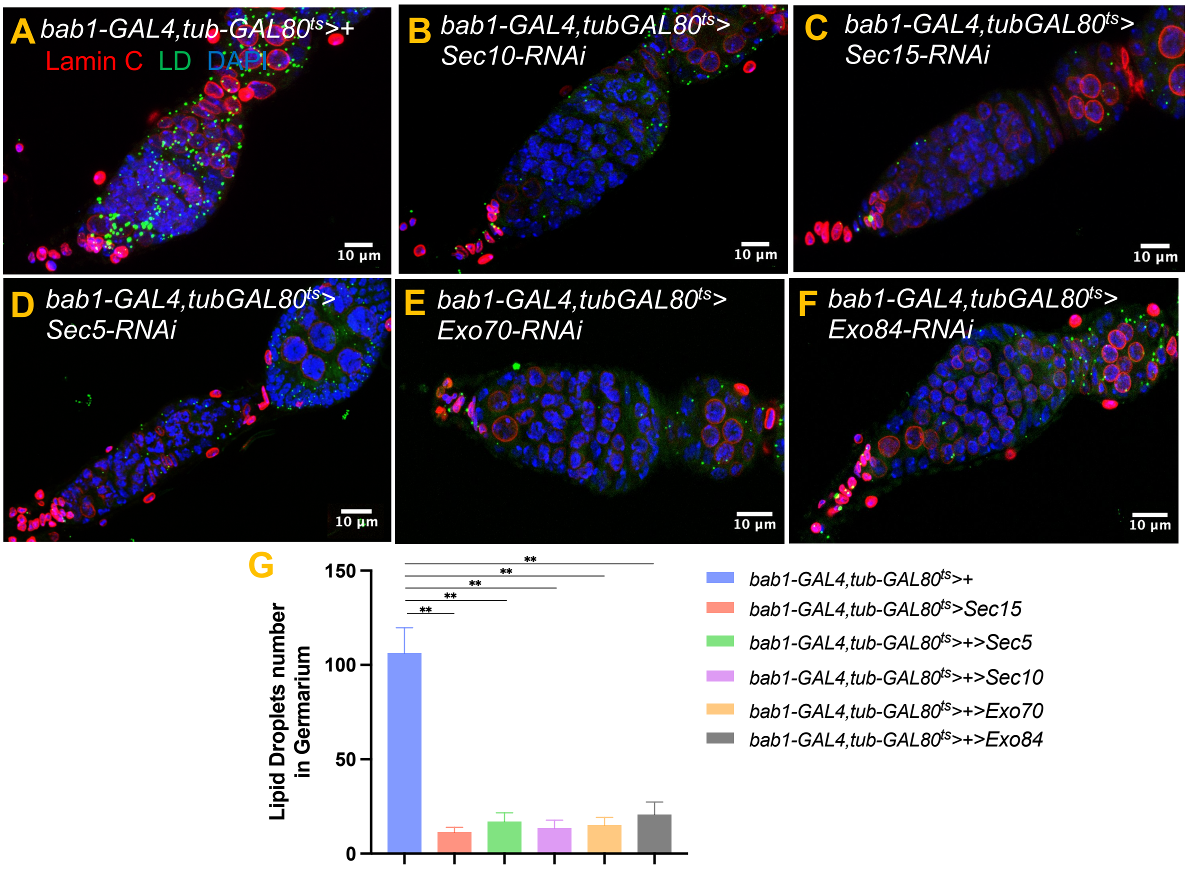


**Figure S3: Exocyst complex component knockdown shows lipid droplet depletion from germarium.**

(**A**)Control showing normal LDs distribution **(B-F)** Conditional knockdown of exocyst genes (*Sec10, Sec15, Sec5, Exo70, Exo84*), showing LD depletion from germ cell of germarium (temperature shifted at day1 of adult eclosion, dissected on day 3). LaminC (TF marker), BODIPY (LDs), and DAPI. (**G**) Quantification of LD in different exocyst component knockdown with reduction in LD throughout the germarium. n=42 represents number of germarium analyzed, statistical significance was determined using unpaired two-tailed t-test, (ns = not significant, *P < 0.05, **P < 0.01, ***P < 0.001).

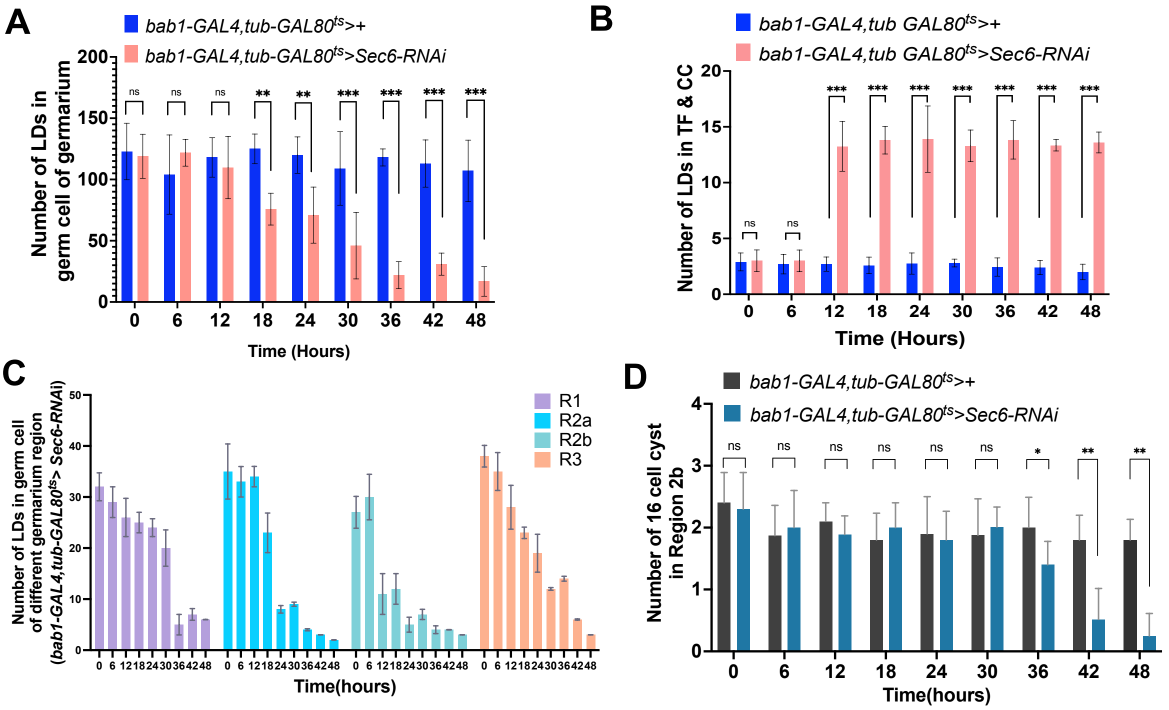


**Figure S4: Sec6-dependent lipid transfer kinetics from TF to germarium.**

**(A)** Lipid Droplet quantification in the germarium showed consistent depletion following *Sec6* knockdown from 18 hours and onwards relative to controls. Data are presented as mean ± SD, n = 52 germarium from each genotype. **(B)** Quantification of LD content in the TF following *Sec6* knockdown confirms significantly elevated lipid levels across time points. Data are presented as mean ± SD, n = 52 germarium from each genotype. **(C)** Graph represents LD count within different regions of germarium in *Sec6* knockdown, were significant number of LD persists in R1 till 24, as compared to R2a, R2b, R3 with sever reduction from 24hours. Data are presented as mean ± SD, n = 52 germarium from each genotype. (**D**) Quantification shows defective 16 cell cyst formation from 36 hours compared to control. Data shown as mean ± SD, n = 48 germarium from each genotype. Statistical significance was determined using unpaired two-tailed t-test, (ns = not significant, *P < 0.05, **P < 0.01, ***P < 0.001, ****P < 0.0001).

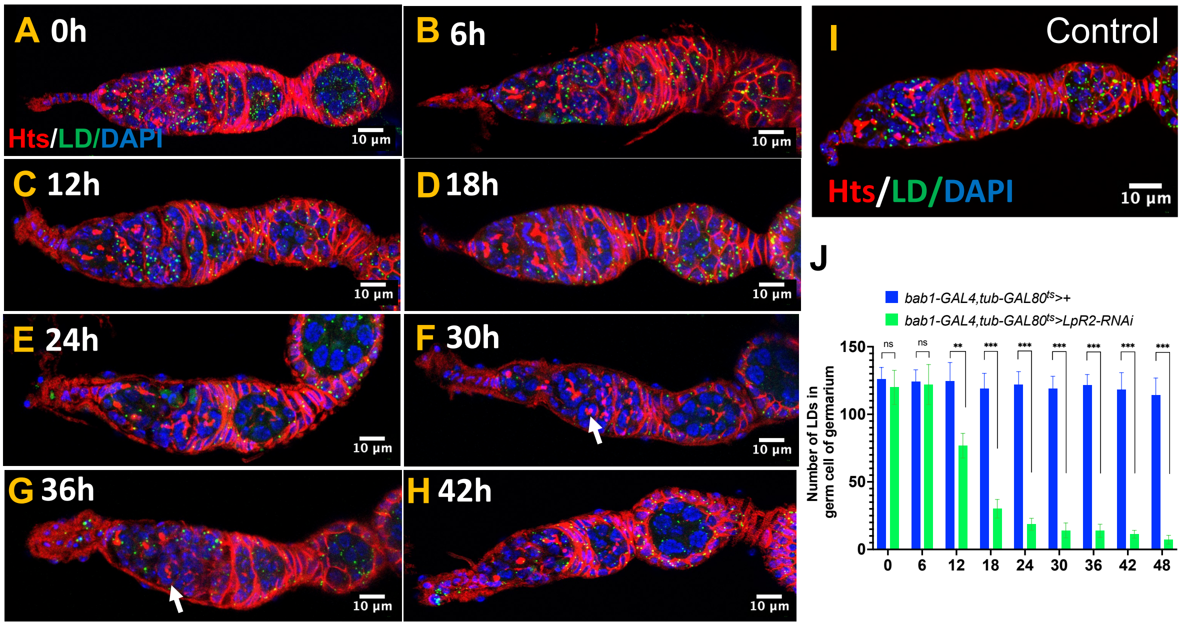


**Figure S5: LpR2 knockdown reduces lipid droplets and causes cyst defects in the germarium.**

**(A-H)** Lipoprotein receptor (*LpR2*) KD in the TF and CC using *bab1-GAL4, tub-GAL80*$^{ts}$ with RNAi induced at 29 °C at day1, and analyzed over 6-hour time interval. LD number depletion within the germarium starts from 12 hour onwards, along with less branched fusome (arrow) and defect in cyst development. **(I)** Control germarium showing lipid droplet distribution and Hts-labeled fusome. **(J)** Quantification of lipid droplets in the germarium reveals a gradual decrease by 12 h, a pronounced reduction from 24 h onward, and a stable, low number at later time points. Data shown as mean ± SD, n = 48 germarium from each genotype. Statistical significance was determined using unpaired two-tailed t-test, (ns = not significant, *P < 0.05, **P < 0.01, ***P < 0.001, ****P < 0.0001).

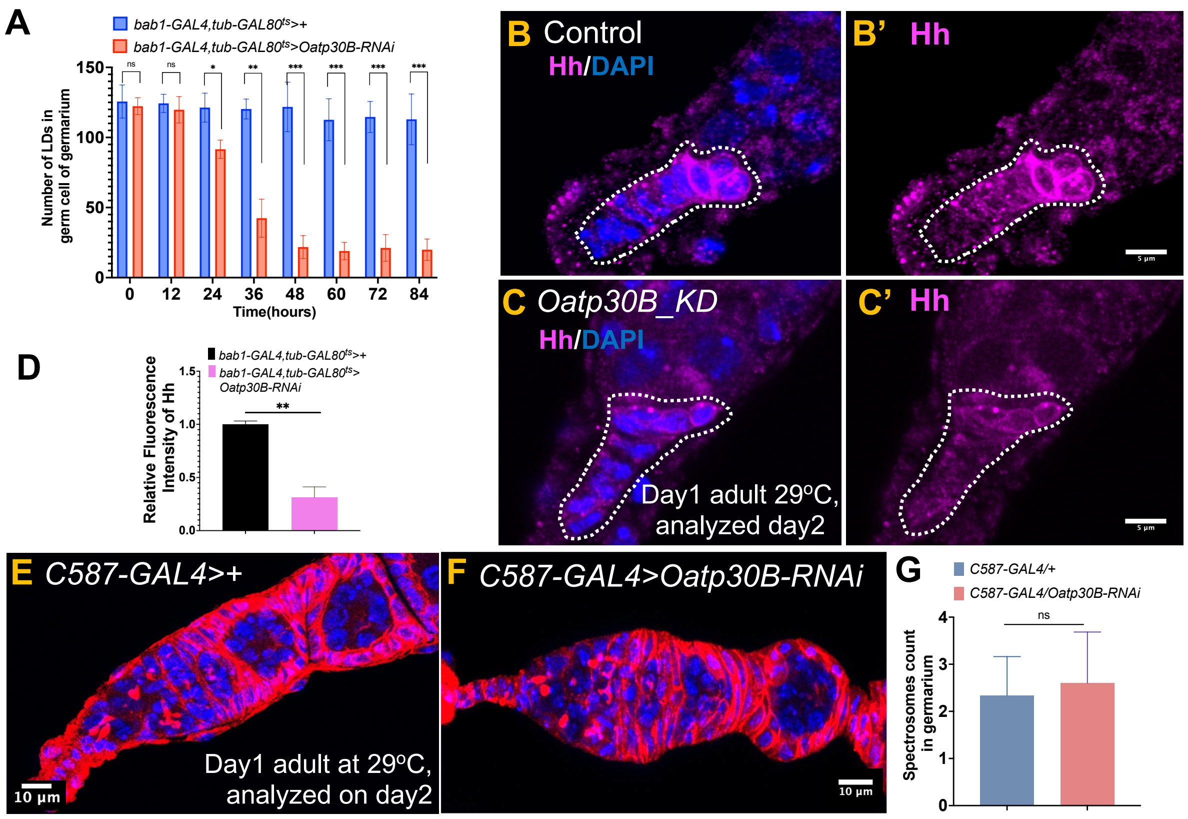


**Figure S6: Terminal filament specific knockdown of *Oatp30B* affect Hedgehog signaling and lipid droplets formation.**

**(A)** Graph representing quantification of lipid droplet numbers in *Oatp30B_KD* germarium over time, showing a decline from 24 to 36 hours, followed by a stable, low number of lipid droplets at later time points. Data shown as mean ± SD, n = 46 germarium analyzed. **(B, B')** Hedgehog (Hh) staining shows robust expression in TF and CC of control germarium. **(C, C')** *Oatp30B* knockdown with *bab1-GAL4, tub-GAL80$^{ts}$* shifted at 29° C at day1 and analyzed at day2, reduced Hh expression in TF and CC cells. **(D)** Quantification of Hh fluorescence intensity in the TF confirms a reduction following *Oatp30B* depletion. Data shown as mean ± SD, n = 35 germarium analyzed. **(E-F)** Representative image of control (E) and escort cell specific *Oatp30B* knockdown (F) germaria driven by c587-GAL4, showing no apparent disruption of cyst or follicle formation. **(G)** Quantification of spectrosome number in *Oatp30B* knockdown and control germarium, showing no significant difference. Data shown as mean ± SD, n = 30 germarium analyzed. Statistical significance was determined using unpaired two-tailed t-test, (ns = not significant, *$P < 0.05$, **$P < 0.01$, ***$P < 0.001$, ****$P < 0.0001$).
